# Data-Driven Virus Discovery with Virushunter and Virusgatherer

**DOI:** 10.64898/2026.01.26.701714

**Authors:** Liubov Chuprikova, Li Chuin Chong, Sergej Ruff, Stefan Seitz, Chris Lauber

## Abstract

We present Virushunter and Virusgatherer, a bipartite computational workflow for discovering viral genomes from unprocessed next-generation sequencing data. Virushunter conducts a profile Hidden Markov Model-based sequence homology search for viral hallmark protein-coding sequence reads to construct microcontigs, which are used as seeds for targeted viral genome assembly by Virusgatherer. We showcase the application of Virushunter and Virusgatherer through an RNA virus-wide screen in 3348 bat transcriptome sequencing experiments from the Sequence Read Archive. Many of the discovered viruses are likely prototypes of new putative virus species, including a novel alphacoronavirus identified in a South American velvety free-tailed bat and a novel rhabdovirus in a European barbastelle bat. The tools are implemented as a reproducible Snakemake pipeline, which is available at https://github.com/lauberlab/VirusHunterGatherer.

## Introduction

Virus discovery has undergone significant advancements in recent years, driven primarily by the massive increase in next-generation sequencing (NGS) data and the parallel development of sophisticated data analysis methods. While traditional virus discovery approaches often relied on sample collection, experimental cultivation, and isolation techniques, data-driven virus discovery (DDVD) (1) leverages vast amounts of publicly accessible NGS data, allowing unprecedented exploration of viral diversity in high-throughput (2–7). DDVD enables researchers to uncover a broad spectrum of viruses, including those with low sequence similarity to known reference viruses. This advancement has opened new avenues for understanding viral evolution, ecology, and virus-host interactions. Several computational tools or resources have been developed to facilitate virus discovery, which do not necessarily follow a DDVD approach, and their descriptions are listed in **Supplementary Table S1**.

Here, we present Virushunter and Virusgatherer, a two-stage computational pipeline that implements a DDVD approach designed to discover both known and divergent novel viruses with high sensitivity from primary sequencing data in a high-throughput manner. The first stage, Virushunter, conducts a profile Hidden Markov Model (pHMM)-based homology search in the raw read data to identify the presence of coding sequences for viral hallmark proteins. Detected viral sequencing reads are assembled to microcontigs that serve as seeds for Virusgatherer, the second stage of the workflow, which performs a targeted assembly of the viral genome sequence. Both user-provided and public sequencing data, e.g. from the Sequence Read Archive (SRA) of the National Center for Biotechnology Information (NCBI), can be used as input for Virushunter and Virusgatherer, and the results are summarized in the form of hit tables and can be visualized using the accompanying R package Virusparies (8). We demonstrate the pipeline through an RNA virus-wide screen in 3,348 publicly available bat transcriptome sequencing experiments from the SRA.

## Material and methods

### Pipeline design and implementation

The Virushunter & Virusgatherer DDVD pipeline is summarized in **Fig. 1**. The main pipeline stages include data acquisition, virus detection through Virushunter, and viral genome assembly through Virusgatherer. Results from Virushunter and Virusgatherer are provided as tab-separated hit tables, which can be visualized using the accompanying R package Virusparies, and as FASTA files containing assembled viral genome sequences. The different pipeline steps are implemented as separate rules, managed by the Snakemake workflow management system (9). All pipeline parameters can be set by the user through the YAML configuration file. The pipeline is optimized for Linux operating systems and depends on several publicly available software tools and reference sequence databases for both hosts and viruses, which can be retrieved from NCBI (see below).

**Figure 1.**
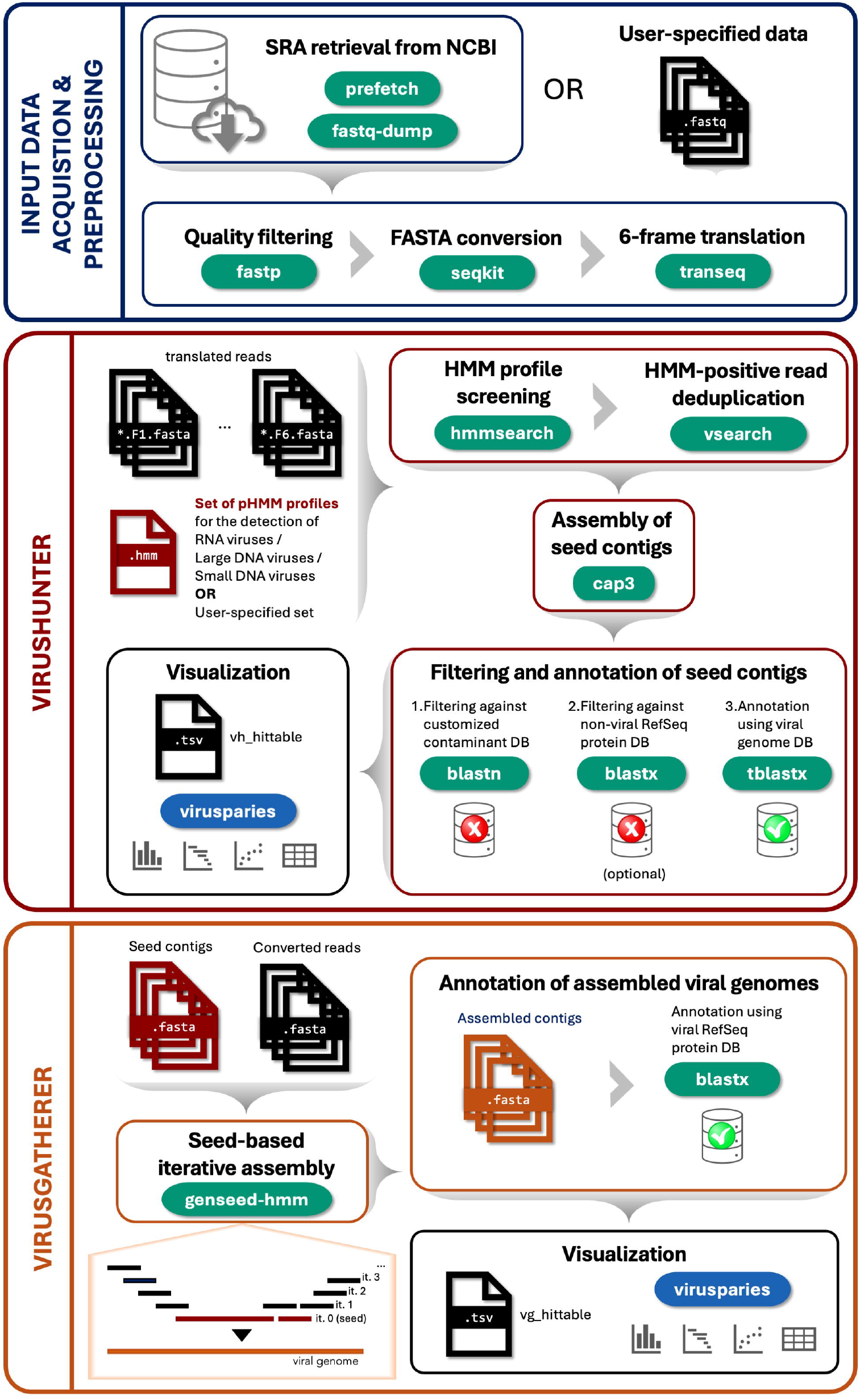
Overview of the Virushunter and Virusgatherer pipeline. The pipeline comprises three main stages: (i) input data acquisition and preprocessing from public (NCBI SRA) or user-provided sequencing data, including quality filtering, conversion to FASTA and translation into six reading frames; (ii) virus detection with Virushunter, where in silico translated reads are screened against viral HMM profiles, deduplicated, assembled into seed contigs, and filtered/annotated against reference databases; and (iii) viral genome reconstruction with Virusgatherer, which performs seed-based iterative assembly and final annotation of assembled viral genomes. Results are provided as FASTA files and tab-separated hit tables that can be visualized using the accompanying Virusparies R package.

The pipeline and installation instructions are available at https://github.com/lauberlab/VirusHunterGatherer. We provide a vhvg.yml file for installing dependencies using the Conda package manager.

### Input data acquisition and preprocessing

Given a list of SRA accession numbers, the pipeline downloads sequencing data from NCBI’s SRA database using prefetch and fastq-dump from the SRA Toolkit (**Fig. 1**, top panel) (10, 11). This step is bypassed if sequencing data in FASTQ format is provided as input by the user. For paired reads, a single interleaved FASTQ file should be generated for each sample. The preprocessing step of the sequencing reads involves trimming sequencing adapters and low-quality bases using fastp (12). The trimmed reads are then converted from FASTQ to FASTA format using seqkit (13) and *in silico* translated into six reading frames using transeq from the EMBOSS suite (14).

### Detecting sequencing reads of viral origin with Virushunter

Virushunter utilizes pHMMs, that were typically constructed at the virus family or higher taxonomic ranks, to screen the raw (unassembled) sequencing data for the presence of reads that share significant sequence similarity to viral marker genes (**Fig. 1**, middle panel). Ideally, these marker genes have no close homologs in cellular genomes to avoid false-positive hits. We provide three sets of pHMMs built on multiple sequence alignments of distinct group-specific key proteins that cover a broad range of (i) RNA viruses, (ii) DNA viruses with small genomes, and (iii) DNA viruses with large genomes (**Supplementary Table S2**). Moreover, users can employ a custom set of pHMMs to screen for any additional virus families (or even mobile genetic elements or cellular genes) of their choice.

The sequence homology search is performed by hmmsearch from the HMMER suite (15) against the 6-frame translated reads, allowing sensitive identification of viral sequences. Reads with an E-value smaller than 10 are deduplicated using vsearch (16) and assembled with CAP3 into microcontigs (can be as large as the employed marker gene) (17). These deduplication and preliminary assembly of the identified viral reads into microcontigs increase the specificity and sensitivity of virus recognition while at the same time reducing the computational time of the subsequent pipeline steps. The resulting microcontigs are filtered by comparing them against a custom database of common contaminants and, optionally, a non-viral reference protein database to remove false-positive hits. The database used in the latter step (such as those provided by the NCBI) has to be considered carefully to avoid contamination with unrecognized viral proteins. The remaining microcontigs are then checked against the NCBI viral reference genome database, retaining hits with an E-value smaller than 1 for the subsequent iterative seed-based assembly by Virusgatherer.

### Assembling viral genome sequences with Virusgatherer

Virusgatherer performs a targeted assembly of viral sequences using Genseed-HMM (18) with the viral microcontigs identified by Virushunter as initial seeds (**Fig. 1**, bottom panel). During iterative contig extension, the ends of the current contig are used as queries to search in blastn mode for additional reads or read pairs aligning to the contig ends. To provisionally annotate the assembled contigs with taxonomic information, they are compared against the NCBI’s database of reference viral proteins (viral RefSeq) via blastx.

Genseed-HMM supports multiple internal assembly programs, and we recommend using CAP3 (17), with Newbler being an alternative (19). Conducting several analyses with different assemblers and combining the results could also be a useful strategy. We distribute a modified version of the original Genseed-HMM code. First, we added a deduplication step after the search for additional reads in each iteration, which significantly decreases CAP3 assembly time for high copy-number viruses. Second, we modified the method for retrieving FASTQ reads due to incompatibilities with recent BLAST versions.

Virusgatherer contigs were clustered using MMSeqs2 with parameters ‘-c 0.5 --cluster-mode 2 --min-seq-id 0.9’ (20), which is not part of the Virusgatherer workflow.

### Reporting and visualizing Virushunter and Virusgatherer results

Both Virushunter and Virusgatherer generate tab-separated hit tables that summarize the results. A Virushunter hit tables provides an initial overview of the viral genomes expected to be present in the sequenced samples, while a Virusgatherer hit table summarizes the viral genome sequences that were assembled from the data. Both hit tables share columns such as the SRA run accession, sample, and study identifiers, along with the host taxon name and taxonomic identifier, all of which are retrieved automatically from the SRA metadata. The Virushunter and Virusgatherer hit tables also provide information about the best match to NCBI’s reference viruses (including subject name, subject taxonomy information, E value, percent amino acid identity) (second sheet in **Supplementary Tables S5 and S6**). Unique to a Virushunter hit table are columns detailing the number of identified reads and (intermediate) microcontigs per input dataset, and the best matching pHMM profile (name and E-value). Columns unique to a Virusgatherer hit table are viral contig name and length.

To facilitate easy inspection and visualization of Virushunter and Virusgatherer results, we provide the accompanying R package **Virusparies** available at CRAN (https://cran.r-project.org/web/packages/Virusparies). Virusparies offers functions to subset and filter hit tables, calculate summary statistics, and generate plots and graphical tables.

## Results

We demonstrate application of the Virushunter and Virusgatherer pipeline through an analysis of 3,348 publicly available transcriptome experiments (as of August 2025) of bat origin obtained from the SRA database (**Supplementary Table S3**). This dataset encompasses sequencing data from 154 species, 79 genera, and 17 families of bats (**Supplementary Table S4**). The SRA run accession list was obtained by querying the SRA website with the taxonomic identifier of the order Chiroptera (taxid 9397) and limiting the result to transcriptomic datasets (source RNA).

### Identification of putative viral sequences using Virushunter

We screened the bat transcriptome experiments for the presence of RNA virus sequences by utilizing a comprehensive collection of 31 RNA virus pHMMs in the Virushunter stage. The pHMMs were constructed at the family level or higher taxonomic ranks and were mostly based on multiple sequence alignments of the hallmark protein of RNA viruses, the RNA-dependent RNA polymerase (RdRp) (**Supplementary Table S2**). Virushunter gave hits in 1,285 (∼38.4%) of the 3,348 analyzed datasets, of which 1,229 produced significant hits (E-value ≤ 1e-5) in a protein BLAST (tblastx) comparison against RNA virus reference sequences. In total, 4,144 unique microcontig hits against different RNA viral reference sequences were obtained, corresponding to 3.2 hits per sequencing experiment on average (**Supplementary Table S5**). The hits were against reference viruses from 74 different RNA virus families but also included false-positive hits against four DNA virus families (**Supplementary Fig. S1**). **Fig. 2A**, produced using the accompanying R package *Virusparies*, illustrates the 20 virus families that ranked top with respect to the number of SRA datasets that were hit. The number of detected sequencing reads of putative viral origin, which is one of the factors for success of viral genome assembly in the subsequent Virusgatherer stage (see below), varied considerably across the different RNA virus families (**Fig. 2B**). For instance, although the number of datasets with *Bornaviridae* sequences was the largest (n=170), only a relatively low number of reads were detected (r=579 in total), suggesting that many of these hits may come from endogenized bornavirus genomes, which are known to be frequent in vertebrates (21). In contrast, *Spinareoviridae* sequences were found in only 43 SRA datasets but showed by far the largest number of detected reads (r=49,201) (**Fig. 2B**), assembling into 91 microcotigs (**Fig. 2C**), suggesting high copy numbers of viral genomes in the positive sequencing experiments which would be expected for actively replicating RNA viruses. The E-values in the filtering Blast analysis against viral reference genomes were in the range from 1e-5 to 1e-173 (**Fig. 2D**) where lower E-values often indicate viral hits highly diversified from the most closely related known reference virus.

**Figure 2.**
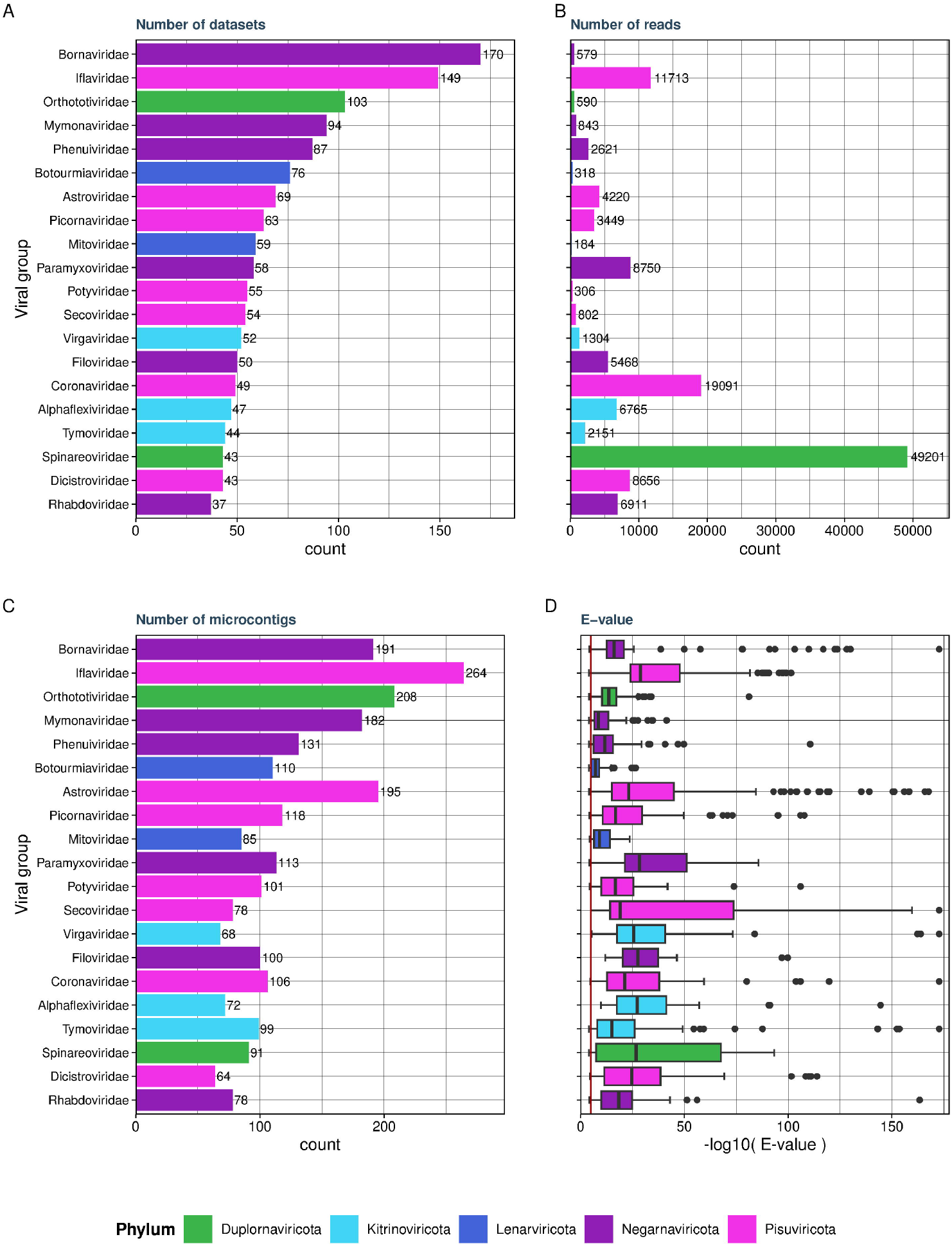
Virushunter results. Results are shown for the 20 RNA virus families with the largest number of SRA datasets in which one or several hits were obtained. Virus families are colored by RNA virus phylum. (A) Number of SRA datasets. (B) Number of sequencing reads summed over all datasets. (C) Number of microcontigs summed over all datasets. (D) Distribution of E-values from protein-based Blast (tblastx) comparison of microcontigs against reference genomes from viral RefSeq. Numbers in A-C indicate absolute counts. All graphs generated using the accompanying R package Virusparies.

Virushunter detects sequence homology to viral marker genes that constitute a small part of the viral genome and therefore provides a first rough overview of the number of discovered putative viruses and their closest known relatives. Viral genome assembly through the Virusgatherer stage is necessary to validate the presence and determine the identity of the viral genome sequences.

### Targeted viral genome sequence assembly using Virusgatherer

Virusgatherer assembled 4,329 contigs ranging from 57 to 27,392 nt in length (**Supplementary Table S6, Supplementary Fig. S2**). To account for potential sequence redundancy, we clustered the contigs at 90% nucleotide sequence identity, resulting in 3,074 contigs, resembling species-like operational taxonomic units (OTUs) with an average length of 497 nt, and 215 (7.0%) and 68 (2.2%) of the contigs being longer than 1,000 and 5,000 nt, respectively. The resulting viral contigs were assigned to 80 RNA virus families with E-values in the range of 1e-5 to 1e-180 or remained unclassified at the family rank and showed protein sequence identities to viral reference sequences between 22 to 100% with an average of 69.6%. Of the 3,074 species-like OTUs, 2441 (79.4%), 1583 (51.5%) and 534 (17.4%) showed sequence identities to viral RefSeq sequences lower than 90%, 70% and 50%, respectively (**Supplementary Table S6, Supplementary Fig. S2**). Among the 20 most abundant RNA virus families were several families known to infect mammals, such as the families *Astroviridae, Paramyxoviridae, Picornaviridae, Coronaviridae, Phenuiviridae*, and *Filoviridae*, but also families whose members are thought to infect invertebrates and even fungi and plants (**Fig. 3**).

**Figure 3.**
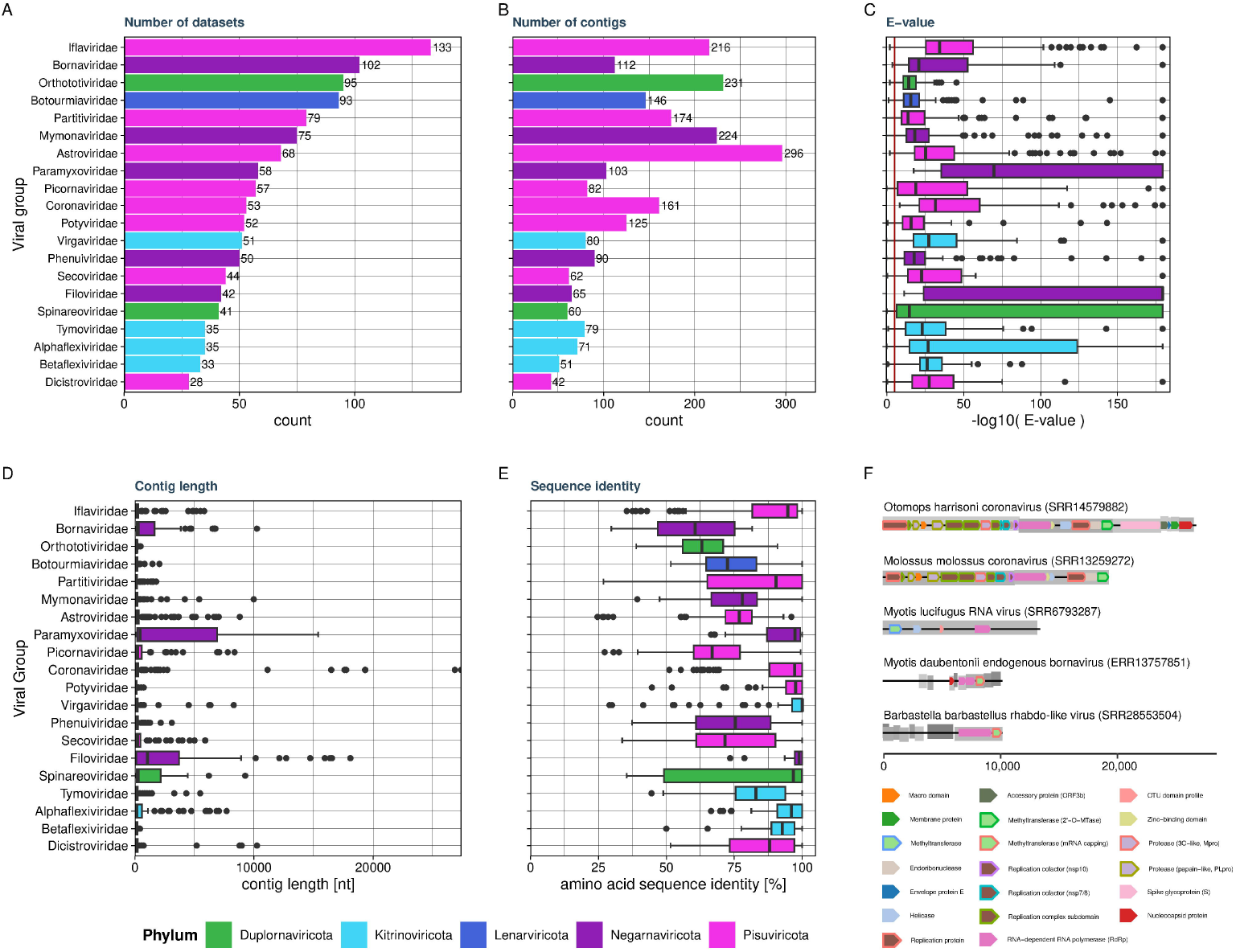
Virusgatherer results. Results are shown for the 20 RNA virus families with the largest number of SRA datasets in which viral contigs were assembled. Virus families are colored by RNA virus phylum. (A) Number of SRA datasets. (B) Number of assembled contigs summed over all datasets. (C) Distribution of E-values from protein-based Blast (tblastx) comparison of contigs against reference genomes from viral RefSeq. (D) Distribution of contig lengths. (E) Distribution of percent amino acid sequence identities from protein-based Blast (tblastx) comparison of contigs against reference genomes from viral RefSeq. (F) Genome maps of five selected viral contigs assembled by Virusgatherer. The contig from SRR13259272 is partial and does not cover the complete protein-coding region of the viral genome. Open reading frames (ORFs) are shown as rectangles with different shadings of gray. Protein domains predicted in an InterProScan analysis are shown as colored shapes within ORFs; see color legend at the bottom. Viruses are named by the bat species in which they were discovered and the most closely related virus reference taxon; corresponding SRA identifiers are in brackets. The scale bar is in nucleotides.

### Discovery of relatives of important human pathogens

Among the discovered RNA viruses were known human pathogens including Zaire ebolavirus, Marburgvirus, and Severe acute respiratory syndrome coronavirus 2 (SARS-CoV-2) (from a project studying the SARS-CoV-2 furin cleavage site in bat cells). We also found viruses closely related to previously reported bat viruses, such as Bat hepacivirus (amino acid sequence identity of 99.97% to accession DAD52839.1), Bat hepatitis E virus (93.73% to XMO98526.1), or Bat morbillivirus (99.89% to DBA64462.1) (**Supplementary Table S6, Supplementary Fig. S3**).

Moreover, we identified a large number of viral genomes with amino acid sequence identities to viral RefSeq entries of well below 90% (**Fig. 3, Supplementary Fig. S2**). Among them was a partial genome sequence covering the ORF1a/b region of a novel coronavirus identified in an intestine sample of a velvety free-tailed bat (species *Molossus molossus*) from Peru (SRA dataset SRR13259272). This novel coronavirus showed an amino acid sequence identity of ∼70.63% to a bat alphacoronavirus (Bat Coronavirus EsJX20, accession WCC63170.1), suggesting that it is a prototype of a new virus species within the genus *Alphacoronavirus*. We also identified a coding-complete genome of a novel rhabdo-like virus in a blood sample of a barbastelle bat (species *Barbastella barbastellus*) from the United Kingdom (SRA dataset SRR28553504) that showed an amino acid sequence identity of only ∼46.65% to an unclassified rhabdovirus (Agrotis ipsilon virus, accession QKY64636.1) (**Fig. 3F, Supplementary Table S6**).

## Discussion

The Virushunter and Virusgatherer DDVD pipeline enables the large-scale screening of raw sequencing data for the presence of viral marker genes and the subsequent reconstruction of respective viral genome sequences. Although we run the presented analysis in a high-performance computing (HPC) environment with 12 to 128 CPUs and a memory of 2 to 4 GB per CPU per SRA experiment, it can be deployed on any modern UNIX-based system.

The pipeline implements a highly sensitive sequence homology search approach based on pHMMs (15). Therefore, it allows for detection of highly divergent viruses (2, 6, 7, 22) that share amino acid sequence identity with reference viruses in the “twilight zone” of only 20-30% (23). In the case of RNA viruses, as analyzed here, the pHMMs primarily target the RdRp. Consequently, only the RdRp-encoding segment of a segmented viral genome will be identified. This shortcoming can be addressed by extending the standard pHMM set with segment-specific pHMMs or using a custom set of pHMMs tailored to particular virus groups.

The degree of “novelty” of an identified virus is based on shared sequence identity to reference viruses and inherently depends on the composition of viral reference genome database employed in Virushunter and Virusgatherer. We therefore suggest regularly updating the locally installed reference sequence databases to the most recent NCBI versions to ensure accurate annotation and novelty assessment.

The success of assembling coding-complete (RNA) viral genomes depends on several factors, including the genetic heterogeneity of the sequenced viral population and the amount of viral genetic material present in the sequenced sample. Genomic regions of high heterogeneity can result in premature termination of the seed-based, iterative assembly process. Standard, untargeted *de novo* assembly could help in some of these cases and may produce longer contigs. The issue of insufficient amounts of viral sequencing reads can only be solved by re-sequencing the sample (6) and is typically due to the fact that the original SRA study was not dedicated to virus discovery. Notwithstanding that, even partial viral genomes provide important insights into previously undescribed viral genetic diversity and viral host range. Their value can be thought of being comparable to that of expressed sequence tags (ESTs) prior to the availability of the human genome sequence (24).

Virushunter and Virusgatherer provide approximate taxonomic assignments of the discovered viral genomes to the most closely related viral taxa recognized by the International Committee on Taxonomy of Viruses (ICTV), but do not provide an accurate taxonomic classification. The clustering based on sequence identity thresholds, as conducted here, which is not part of the Virushunter and Virusgatherer workflow, offers a coarse estimation of the number of viral OTUs. We suggest employing coherent approaches and dedicated tools for the non-trivial and often virus family-specific task of virus taxonomy (25, 26).

## Data availability

The Virushunter and Virusgatherer pipeline is available from GitHub at https://github.com/lauberlab/VirusHunterGatherer. Assembled sequence data accompanying the study have been uploaded to FigShare: DOI 10.6084/m9.figshare.31149460.

## Supplementary data

**Supplementary Figure S1**. Full Virushunter results grouped by virus family and colored by virus phylum. (A) Number of identified SRA datasets, (B) number of identified sequencing reads, (C) number of microcontigs, (D) distribution of E-values of protein Blast search against RNA virus reference sequences.

**Supplementary Figure S2**. Full Virusgatherer results grouped by virus family and colored by virus phylum. (A) Number of identified SRA datasets, (B) number of contigs, (C) distribution of E-values of protein Blast search against RNA virus reference sequences, (D) distribution of contig lengths, (E) distribution of protein sequence identity to closest reference.

**Supplementary Figure S3**. Genome maps of 33 discovered non-redundant RNA virus genomes of at least 5000 nt and amino acid sequence identity to viral RefSeq representatives of less than 80%. Open reading frames (ORFs) are shown as rectangles with different shadings of gray. Protein domains predicted in an InterProScan analysis are shown as colored shapes within ORFs; see color legend at the bottom. For detailed virus annotation see Supplementary Table S6.

**Supplementary Table S1**. Computational virus discovery tools.

**Supplementary Table S2**. Profile Hidden Markov Models used in the Virushunter search and their taxonomic coverage.

**Supplementary Table S3**. List of bat transcriptome sequencing runs publicly available at the SRA as of August 2025.

**Supplementary Table S4**. Taxonomic annotation associated with the analyzed bat transcriptome sequencing runs.

**Supplementary Table S5**. Virushunter hit table for search in 3,348 bat SRA transcriptome experiments (sheet 1) and description of columns (sheet 2).

**Supplementary Table S6**. Virusgatherer hit table for search in 3,348 bat SRA transcriptome experiments (sheet 1) and description of columns (sheet 2).

## Acknowledgements

We thank all colleagues in the scientific community who make their sequence data publicly accessible. We acknowledge the NCBI for providing an elaborate platform to exchange sequencing data. We gratefully acknowledge the computing time made available to us on the high-performance computers Romeo and Barnard at the NHR (Nationales Hochleistungsrechnen an Hochschulen) Center NHR@TUD of the University of Technology Dresden.

## Author contributions

L.C. (Formal analysis, Software, Visualization, Writing – original draft), L.C.C. (Formal analysis, Software, Visualization, Writing – review & editing), S.R. (Formal analysis, Software, Visualization, Writing – review & editing), S.S. (Conceptualization, Funding acquisition, Supervision, Writing – review & editing), C.L. (Conceptualization, Formal analysis, Funding acquisition, Software, Supervision, Visualization, Writing – original draft).

