## Supplementary figures and images for "Data-Driven Virus Discovery with Virushunter and Virusgatherer"

### Supplementary Figure 1

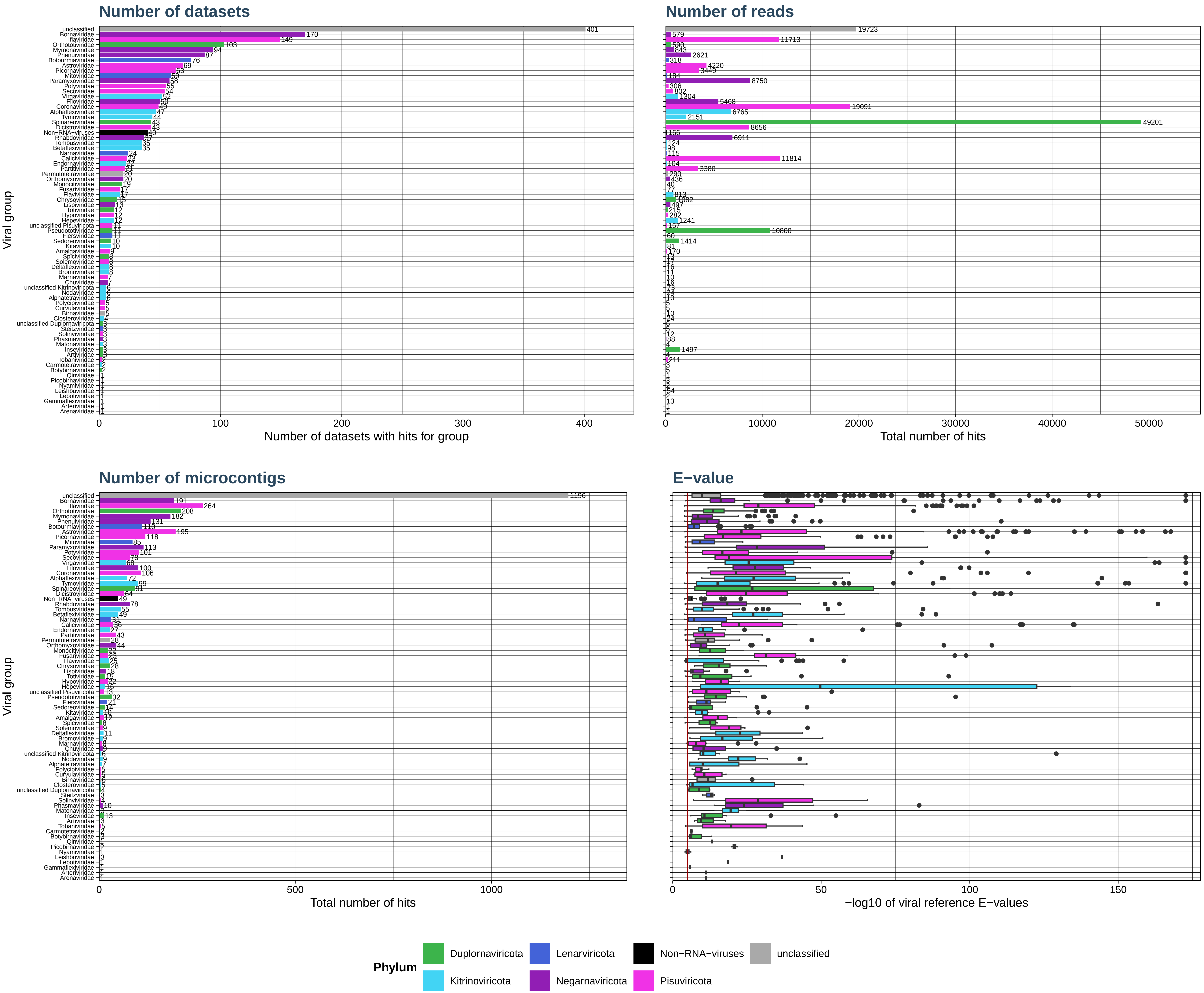

### Supplementary Figure 2

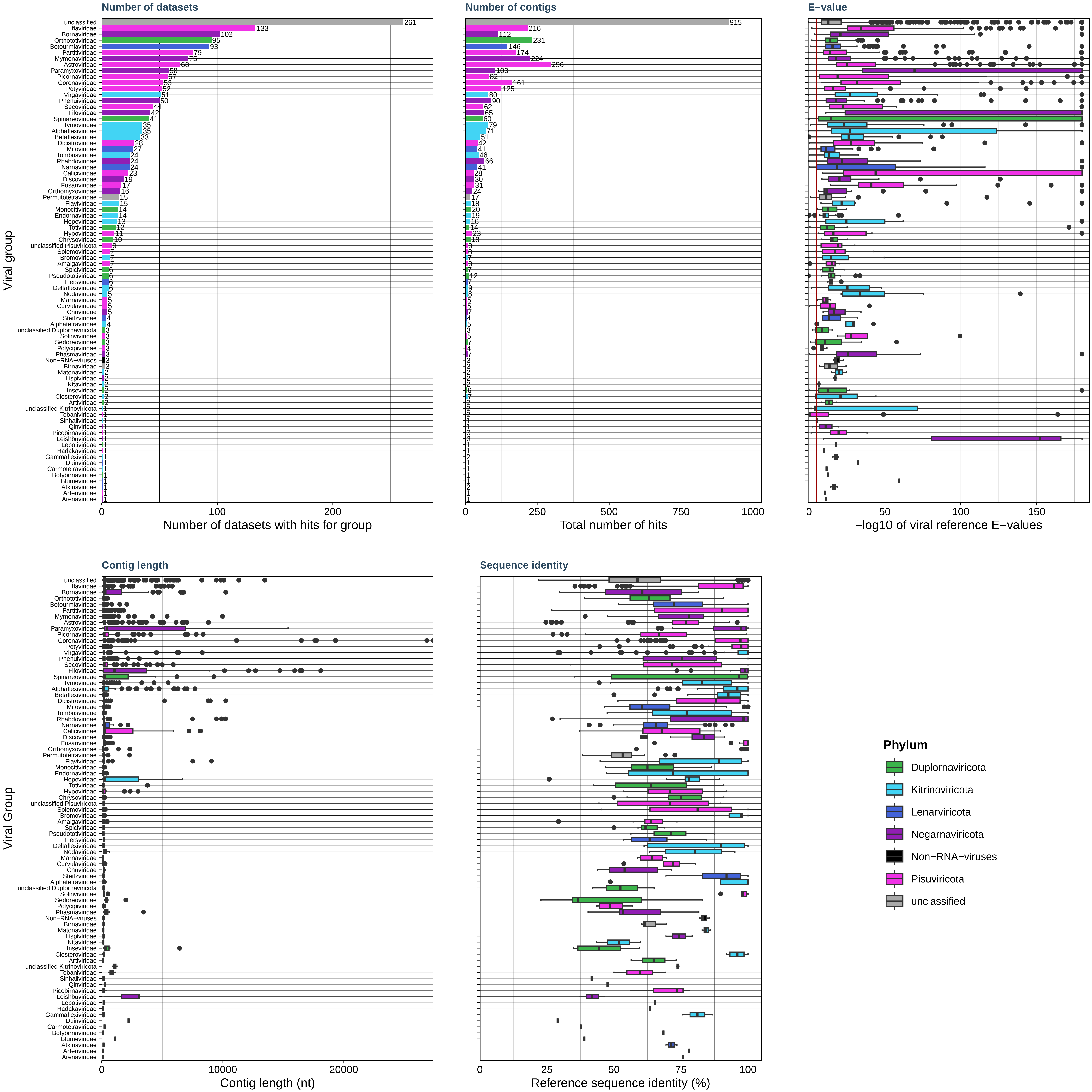

### Supplementary Figure 3

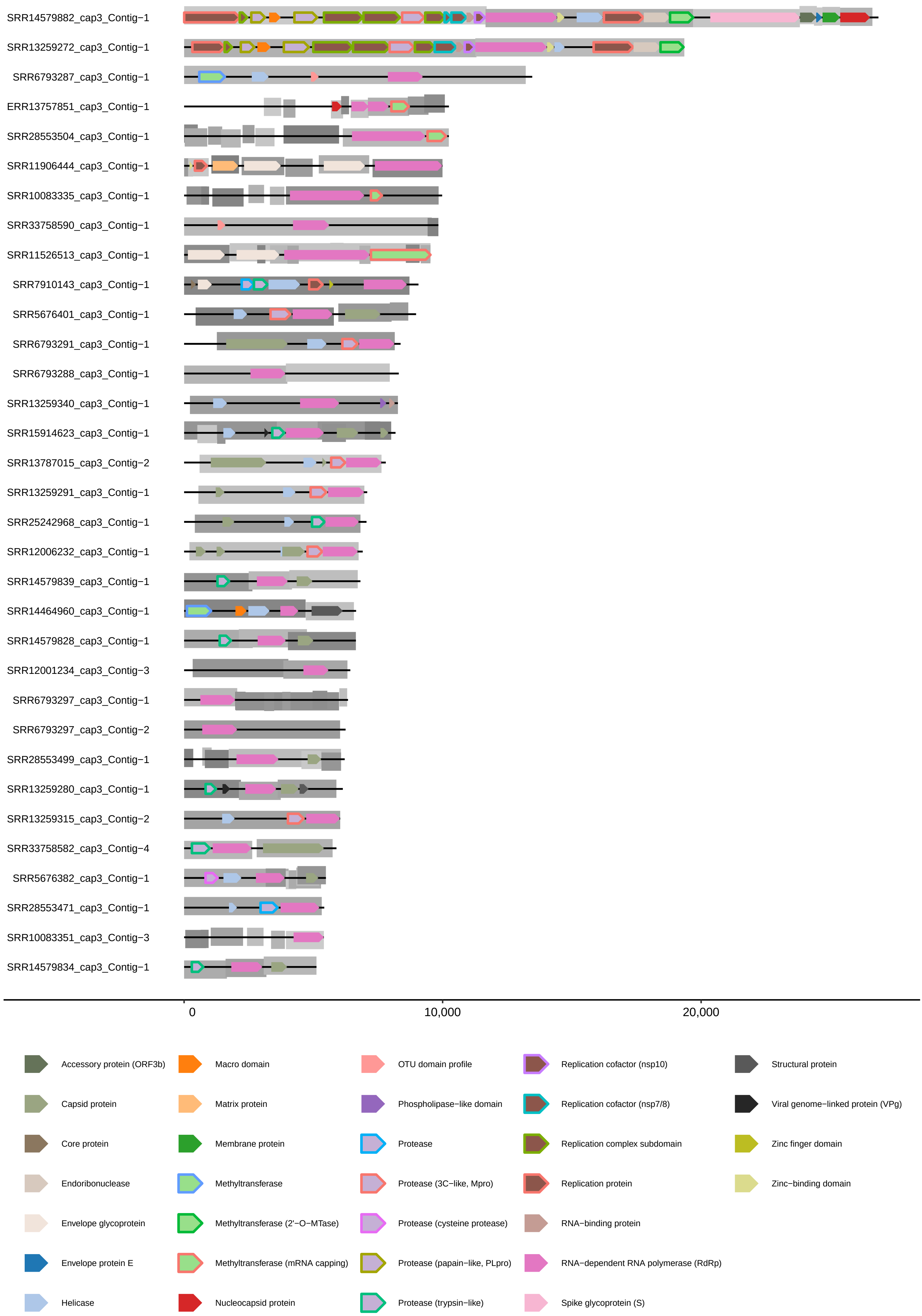
